# CelLEVITAS: Label-free rapid sorting and enrichment of live cells via magnetic levitation

**DOI:** 10.1101/2020.07.27.223917

**Authors:** Elliot K. Chin, Colin A. Grant, Mehmet Giray Ogut, Bocheng Cai, Naside Gozde Durmus

## Abstract

Sorting methods that remove non-viable cells and debris, while retaining a high yield of viable cells, are crucial for many applications in biotechnology, genomics, tissue engineering and medicine. However, a significant challenge is gentle sorting of these different cell states based on very minute differences in density and magnetic signatures, without relying on any labels, tags or markers. Here, a new magnetic levitation-based technology, CelLEVITAS, is developed for the label-free sorting and enrichment of live cells. This work reports the first use of magnetic levitation for sorting of viable and non-viable cells within a microfluidic device and demonstrates extremely effective removal of dead cells and debris from heterogeneous samples. First, the levitation conditions for separating viable and non-viable cells under a magnetic field were fine-tuned. Levitation trajectories of live and dead cell states were then monitored in real-time, as cells magnetically focused to their corresponding levitation bands. CelLEVITAS successfully sorted and enriched live cells from a variety of input cell concentrations (100-200,000 cells/mL) and a variety of input purities (10-50%) into consistently high output purities (>80%). This method is sensitive, does not impair cell viability during sorting, and significantly increases the input sample viability up to 7-fold. Overall, this new magnetic levitation-based sorting strategy drastically reduces the processing time to a single-step, 30-minute sorting protocol and eliminates the manual pre-processing and labeling steps that are required for traditional flow cytometry techniques.

## 1. Introduction

Sorting cells is of great interest in biotechnology, regenerative medicine, drug screening, tissue engineering, and bio printing. Specifically, differentiating and separating viable and non-viable cells is vital in many fields. For instance, separating live cells has many important applications in medicine, especially for drug screening and toxicology studies. The ability to separate and isolate non-viable cells from a living population is crucial for predicting the toxicity of drug compounds, saving considerable time and money in the drug discovery pipeline.^[1]^ Among other fundamental applications of live-dead separation are cell-based therapies such as hematopoietic stem cell transplants, where the capability to separate apoptotic cells bears serious consequence for the success of stem cell grafts.^[2]^ Other uses of stem cells, such as for bio printing, also are inhibited by low cell viability, because insufficient viability skews differentiation of stem cells.^[3,4]^ Furthermore, removing dead cells inside a cell culture is pivotal in general cell culture maintenance. In biotechnology, increasing the cell viability of bioreactors improves antibody secretion.^[5,6]^ Thus, from assessing viability of cell-cultures in a laboratory to targeted induction of apoptosis during neurological regenerative therapy, live-dead cell sorting is an essential and common procedure.^[7]^

Separating viable and non-viable cells is usually tedious and painstaking with conventional label-based sorting mechanisms, such as fluorescence microscopy, flow cytometry and microplate readers. This is because applications that rely on staining are prone to high entry costs, negative impacts on the lifecycle of affected cells, and false positive signals due to the nonspecific binding of reagents.^[8–10]^ Furthermore, magnetic nanoparticles, commonly used in label-based magnetic-activated cell sorting (MACS), are toxic to certain cell types.^[11]^ In addition, centrifugation might damage sensitive cells, such as leukocytes.^[10,12,13]^ Generally, the most commonly used sorting methods are cumbersome due to the pre-requisite of many labels. Many label-free microfluidic techniques have thus been developed to isolate live cells using physical attributes including size, density, stiffness and electrical polarizability. For example, microscale filtration technologies select for live cells based on size and deformability by passing a solution through various forms of filters.^[10]^ However, these filters are prone to clogging, resulting in turbulence, and fail to account for frequent variation in cell sizes.^[10]^ Another size-based method is acoustophoresis, where acoustic pressure waves move larger live cells to a selection channel, removing dead cells that have shrunk as part of the apoptotic mechanism.^[10]^ Acoustophoretic channels are easily clogged, and increasing flow rate and acoustic pressure to compensate can damage cells.^[14,15]^ An alternative sorting method that leverages the dielectric properties of cells for live sorting is dielectrophoresis.^[16]^ Although dielectrophoresis can differentiate between live and dead cells, its throughput is low at no more than .36 mL/hour.^[10]^ These techniques are cost effective and do not suffer from high false positives as described for label dependent sorting technologies. Furthermore, they are often more cost-effective and automatable. However, because these current label-free techniques rely solely on external physical attributes, they can lack selectivity. Therefore, there is a need for new inexpensive, gentle and efficient label-free sorting methods.^[10]^

Here, we report the first use of magnetic levitation for label-free sorting of viable and non-viable cells within a microfluidic device, and demonstrate extremely effective removal of dead cells and debris from heterogeneous samples, even with very low input cell viability (10%). Although we demonstrated earlier that cells can be levitated to varying heights under a magnetic field, we have not yet presented the application of magnetic levitation to sort cells in live and dead states by their viability. In this work, we further created a new microfluidic sorting device that exploits the differences in density and magnetic susceptibility between viable and non-viable cell states. We name this new device CelLEVITAS. First, we fine-tuned and optimized the paramagnetic medium conditions for levitating and separating viable and non-viable cells under a magnetic field. In addition, we showed that the levitation trajectories of live and dead cell states can be monitored in real-time, as levitated cells were magnetically focused to their corresponding levitation bands. Leveraging these discoveries, viable breast cancer cells were selectively enriched from heterogeneous mixtures of live and dead cells. We showed that the CelLEVITAS device can successfully sort and enrich live cells from a variety of input cell concentrations (100-200,000 cells/mL) and a variety of input purities (10-50%) into consistently high output purities (>80%). CelLEVITAS is sensitive and it does not impair cell viability during sorting, and significantly increases the input sample viability up to 7-fold. Overall, this new magnetic levitation-based separation strategy drastically reduces the processing time of cell sorting to a single-step, 30-minute protocol and eliminates the manual pre-processing steps, such as centrifugation, pre-labelling and pre-staining, that are usually required for traditional flow cytometry techniques **(Figure S1)**. It is envisioned that this inexpensive, label-free, gentle sorting method can be broadly applicable for many cell types with heteregenous cell states.

## 2. Results

### 2.1 Separation of Cells Based on the Principles of Magnetic Levitation

CelLEVITAS is a densitometry and imaging platform which uses the principles of magnetic levitation. CelLEVITAS device consists of a flow channel (1 mm in height and width) held between two custom-designed rare earth magnets for gentle and rapid separation of different cell states in a continuous flow operation. The system is driven by two syringe pumps that withdraw the sample into the flow-based magnetic levitation system. For real-time imaging, aluminum-coated mirrors are placed at each side of the microchannel and a camera images the cells as they levitate and move through the flow channel. In this unique device configuration, cells can be levitated and separated based on their inherent density without the use of labels, antibodies or tags.^[1]^ The magnetic susceptibility difference between a cell and its surrounding paramagnetic medium causes it to move away from a higher (*i.e*., close vicinity at the magnets) to a lower magnetic field strength site (*i.e*., away from the magnets) until gravitational, buoyancy and magnetic forces acting on the cells reach an equilibrium. Cells are levitated at a final position between the two magnets, where the magnetic force (F_mag_) equals the buoyancy force (F_b_). As we have reported earlier, the equilibrium height of a levitating cell mainly depends on its inherent density signature.^[1]^ Here, we further applied magnetic levitation profiling to separate and enrich cells based on their biological states (i.e., live, viable cells), according to density differences **(Figure 1)**. A heterogeneous sample containing the live and dead/dying cell populations are loaded in a non-ionic paramagnetic medium and introduced into the levitation-based sorting platform (**Step 1**). As the cells flow, the platform monitors the equilibrium heights of each individual cell in real-time, magnetically focuses and separates live and dead/dying cells at different levitation bands, based on their unique magnetic and density signatures **(Step 2)**. After separation, live cells (lighter portion) can be sorted to the top outlet, while the dead/dying cells and debris (denser portion) are collected to the bottom outlet (**Step 3**). In essence, the system can act as a “single-cell centrifuge” through the magnet-induced density gradients to precisely detect and separate different cell types and physiological states. Overall, this new magnetic levitation-based sorting strategy drastically reduces the processing time to a single-step, 30-minute sorting protocol and eliminates the manual pre-processing steps, such as centrifugation and pre-staining, that are usually required for traditional cell sorting technologies.

**Figure 1.**
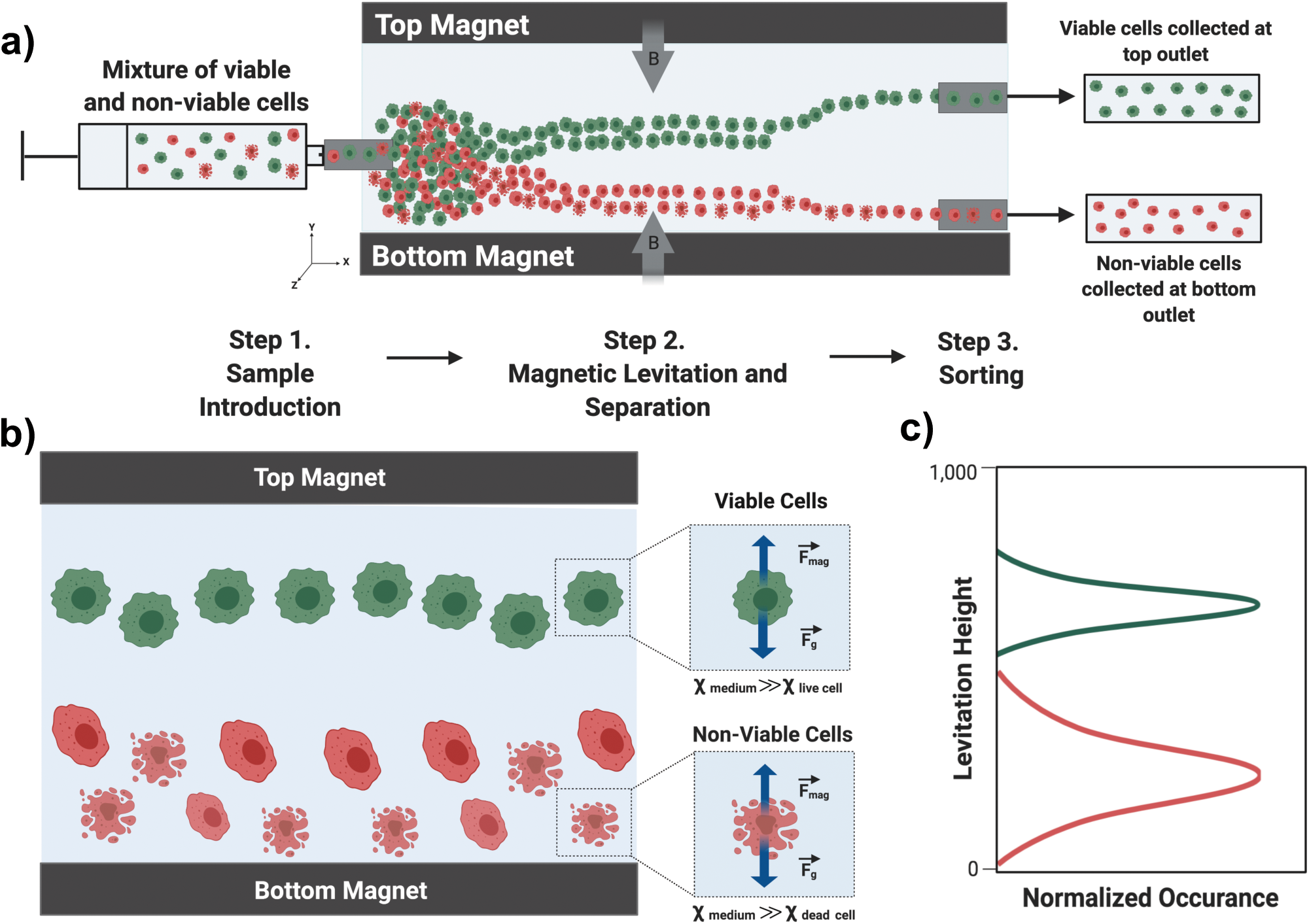
Label-free, high-throughput sorting and enrichment of live cells via magnetic levitation. **a)** CelLEVITAS uses the principles of magnetic levitation to achieve label-free, high throughput separation of live and non-viable cells. A heterogeneous sample containing the live (green) and dead/dying (red) cell populations are loaded in a paramagnetic medium and introduced into the inlet of levitation sorting platform (**Step 1**). CelLEVITAS magnetically focuses and separates the live and dead/dying cells at different levitation bands, based on their unique magnetic and density signatures. As the sample flows through the system, live and dead cells can be imaged, and counted (**Step 2**). After separation, live cells (lighter portion) are sorted to the top outlet, while the dead/dying cells and debris (denser portion) are collected to the bottom outlet (**Step 3**). **b)** The system acts as a “single-cell centrifuge” through the magnet-induced density gradients to precisely detect and separate cells with different physiological states. Cells levitate where their magnetic and gravitational forces are equal to their buoyancy force, determined by their position in the density gradient generated within the static device. **c)** Viable and nonviable cells, due to different densities and magnetic susceptibilities, levitate on different equilibrium planes. Overall, this magnetic levitation-based sorting strategy drastically reduces the sample processing time to a single-step, 30 minutes protocol and eliminates the manual pre-processing steps, such as centrifugation, pre-staining or pre-labelling, that are commonly required for traditional cell sorting technologies.

### 2.2 CelLEVITAS Levitation of Live and Dead Cell States

While cell death occurs, cells change in size, surface-to-volume ratio, and density.^[2]^ As a proof-of-concept, we first investigated the levitation characteristics of live and dead cells in the static (non-flow) magnetic levitation system **(Figure 2)**. For our experiments, we used heterogeneous mixtures of viable and non-viable MDA-MB-231 breast cancer cells, a cell line widely used in cancer research.^[17]^ For identification and confirmation of different cell states of interest, live MDA-MB-231 cells were stained with Calcein-AM (green) and dead MDA-MB-231 cells were stained with propidium iodide (red). Then, live and dead sub-populations were mixed and levitated at 30 mM paramagnetic medium. Within 15 minutes, live and dead/dying cells equilibrated at different heights. Live cells maintained their equilibrium height and formed a distinct band, while denser dead/dying cells sink to the bottom of the microchannel **(Figure 2a)**. The levitation heights of live and dead cell states were fine-tuned at various paramagnetic medium concentrations (i.e., 50 mM and 100 mM). Increasing the paramagnetic medium concentration enabled us to focus the dead cells at higher heights. In addition, using fluorescent markers as a measure of viability, we showed that live and dead cells formed two distinct bands at all the paramagnetic concentrations tested. Then, levitation heights were analyzed using an in-house developed image analysis software that the finds the positions of live and dead cells using an image recognition algorithm **(Figure S2)**. Using this algorithm, we quantified and optimized the separation distance between live and dead cell states at different paramagnetic concentrations (i.e., 30 mM, 50 mM, and 100 mM) **(Figure 2b)**. At 30 mM, live cells had an average levitation height of 233.35 ± 17.03 µm, while dead/dying cells had an average height of 65.20 ± 19.19 µm. This resulted in a 168.16 ± 33.71 µm separation distance between live and dead populations **(Figure 2c)**. At 50 mM, live cells had an average levitation height of 351.74 ± 18.04 µm, while dead/dying cells had an average levitation height of 115.84 ± 30.41 µm. This resulted in a 235.90 ± 13.39 µm separation distance. At 100 mM, live cells had an average levitation height of 362.33 ± 22.23 µm, while dead/dying cells had an average levitation height of 195.72 ± 8.93 µm. This resulted in a 169.64 ± 17.17 µm separation distance. Thus, 50 mM paramagnetic medium resulted in the best separation distance between the live and dead/dying cells in the levitation system.

**Figure 2.**
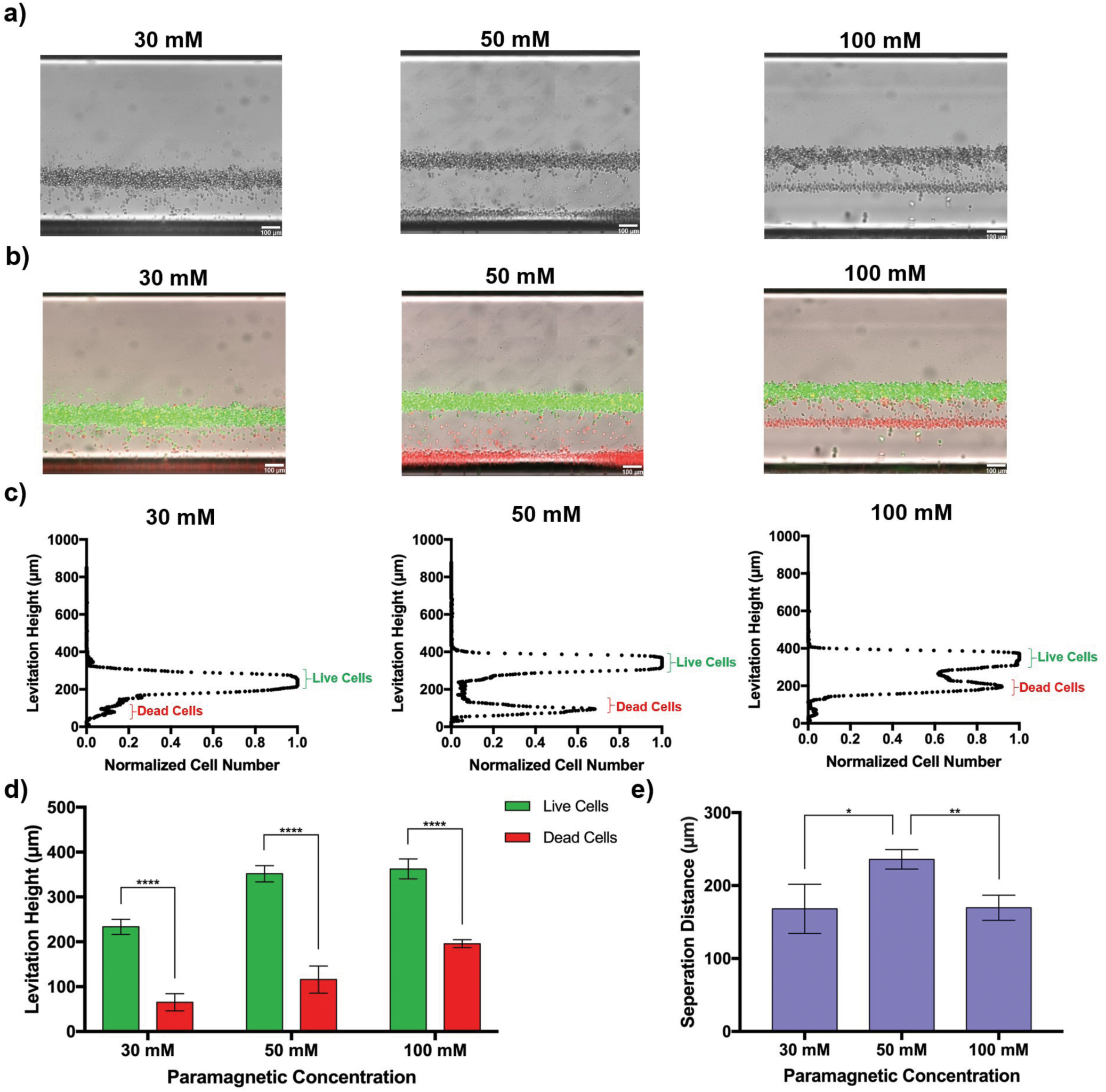
Magnetic levitation and separation of live and dead cell states. **a)** Live and dead sub-populations of MDA-MB-231 breast cancer cells were mixed and levitated at different paramagnetic medium concentrations (i.e., 30 mM, 50 mM and 100 mM). Live and dead/dying cells formed two distinct levitation bands at all the paramagnetic concentrations tested. **b)** Fluorescent imaging of these bands, where live cells are stained with calcine (green) and dead cells with propidium iodide (red), displays clear separation between viable and nonviable cells. **c, d)** Analysis and quantification of levitation heights of live and dead cells at different paramagnetic medium concentrations. Live cells levitate higher than the dead/dying cells. **e)** Optimization of the separation distance between different cell states. 50 mM paramagnetic medium yielded the best separation distance between the live and dead/dying cells in the static levitation system. * p < .05; ** p<.01; *** p<.001; **** p<.0001, by unequal variances t-test.

### 2.3 Real-time Monitoring and Magnetic Focusing of Live and Dead Cells

Next, we monitored the magnetic focusing of live and dead cells and quantified their heterogeneous responses in real-time **(Figure 3)**. A mixture of live and dead cancer cells was levitated in 50 mM paramagnetic medium for 30 minutes and imaged every minute. Notably, live cells (green) focused at their unique levitation band within minutes, while dead/dying cells (red) slowly settled down to the bottom of the channel **(Figure 3a and b)**. Next, levitation trajectories of live and dead/dying cells were analyzed and quantified over time **(Figure 3c)**. Levitation profiles of live and dead/dying cell populations were analyzed and quantified by an in-house developed image analysis software **(Figure S2)**. According to our image analysis, magnetic focusing of live cells occurred very rapidly within 10 minutes. Live cells maintained their average levitation height around 367.14 µm during levitation in 50 mM paramagnetic medium over 30 minutes. On the other hand, dynamic changes and shifts in levitation heights of dead cells were observed, as the dead cells slowly settled to the bottom of the channel, as their average levitation height decreased from 740 µm to 58 µm over time. In addition, the heterogeneity of live and dead cell populations decreased over time, as they equilibrated and magnetically focused to their respective levitation bands **(Figure S3)**. Thus, our analysis showed that characteristic cellular levitation profiles change dynamically in response to the perturbations in the biological states, leading to magnetic focusing and separation of live and dead cells within the levitation system.

**Figure 3.**
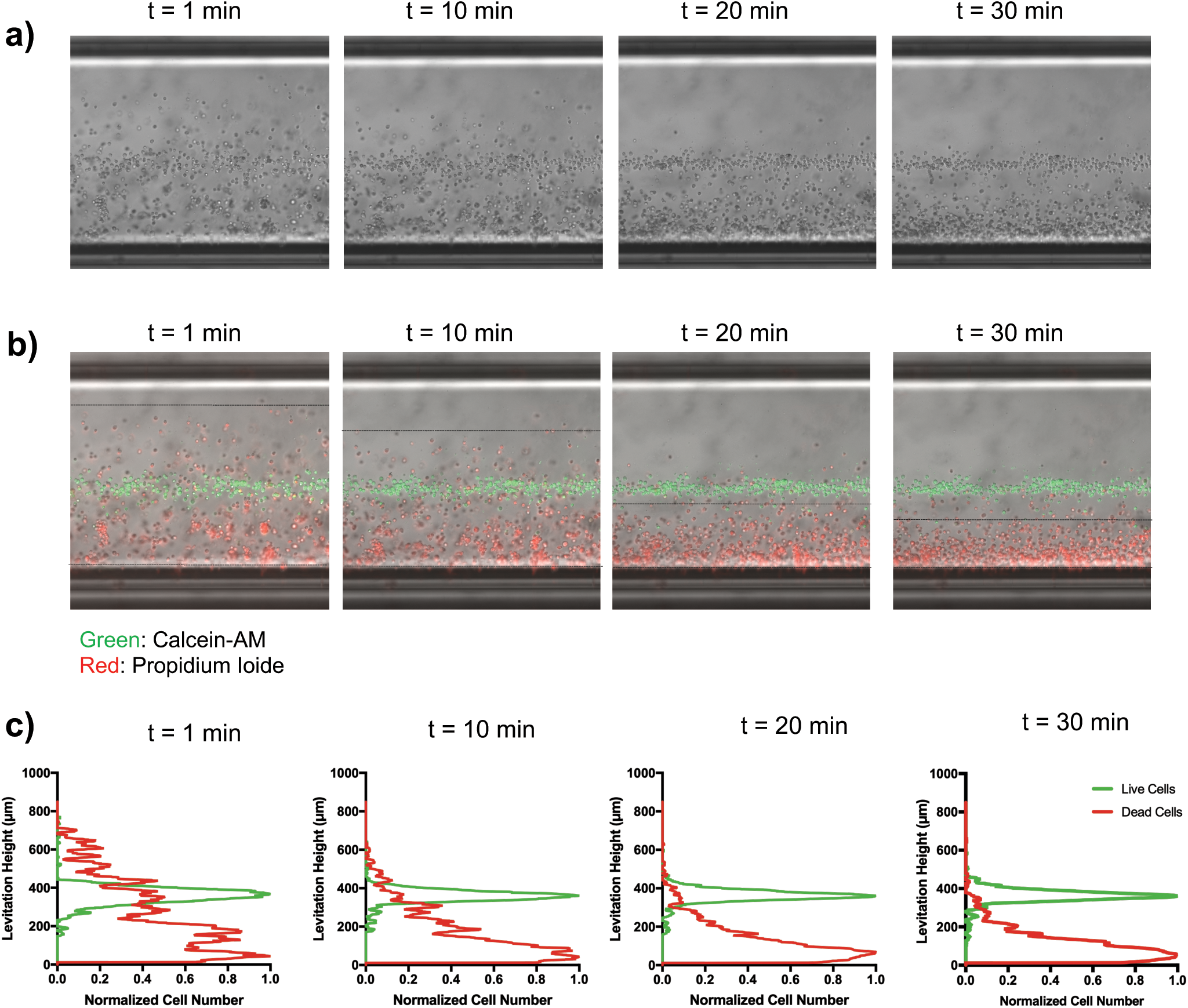
Real-time monitoring and magnetic focusing of live cells from dead cells. **a, b)** Live and dead cell mixture was levitated in 50 mM paramagnetic medium for 30 minutes and imaged every minute. Live cells (green) focused at their unique levitation band within minutes, while dead/dying cells (red) slowly settled down to the bottom of the channel. Gradual decrease in the levitation heights of dead/dying cells is shown with the black lines. Live (green) and dead/dying cell (red) bands are shown in the final image. **c)** Levitation trajectories of live and dead/dying cells over time. Levitation profiles of live and dead/dying cell populations were analyzed and quantified by an in-house developed image analysis software.

### 2.4 Sorting of Live and Dead cells via Magnetic Levitation

To further demonstrate the application of the magnetic levitation method for the enrichment of live cells, we integrated microfluidics to sort the two levitated bands of cells and increase the throughput of the system (**Figure 4**). Heterogeneously mixed live and dead cells, with total concentration of 50-100K cells/mL, were suspended in 50 mM paramagnetic medium and introduced into the CelLEVITAS platform. The mixture was then processed by withdrawing the sample from two outlet channels. Tunable flow rates at the top and bottom outlets enabled precise control and magnetic manipulation of samples along the microfluidic channel. Live cells (stained green) were collected at the top outlet, while dead cells (stained red) were collected at the bottom outlet, as shown in **Figure 4a**. This magnetic levitation-based sorting method is flexible and versatile. It can sort and enrich live cells from very low input cell concentrations (i.e., 100-500 cells/mL) **(Figure 4a, left panel)**, as well as from heterogeneous cell populations from higher input cell concentrations (i.e., >200,000 cells/mL) with higher sample volumes (i.e., 1-10 mL) **(Figure 4a, right panel)**. Heterogeneous input, enriched live cells at the top outlet, and dead cells/debris collected at the bottom collection outlet are shown in **Figure 4b**. Samples with different input purities ranging from 50% to 10%, were processed. In all these different sorting runs, a dramatic removal of dead cells and cell debris was demonstrated (**Figure 4c-d**). Our sorting results indicates that the top outlet achieved high cell viability from as low as 10% input purity, indicating successful sorting and enrichment of live cells, independent of the input purity. This experiment was performed multiple times (n=5), with aggregate results shown in **Fig. 4d. Fig. 4c** highlights three specific trials, where 50% viable cells were purified to 74% viability, 30% viable cells were purified to 75% viability, and 10% viable cells were purified to 76% viability, respectively. Across all trials, CelLEVITAS sorting created a statistically different (p=.0028) population by unequal variance t-test (n=5). Given the levitation information collected in the static system, separation of live cancer cells from dead cancer cells and debris was performed using a total flow rate of 1.2 mL/hour and a 2:1 top: bottom flow rate ratio. Recovery values indicated that over 80% of viable cells were recovered at the top outlet, as shown in **Figure S4**. Importantly, the cells sorted through the system preserved their viability. In summary, CelLEVITAS is a gentle method that can purify samples with a wide range of starting viability into a consistently high viability output, while retaining a high viable cell recovery rate and throughput.

**Figure 4.**
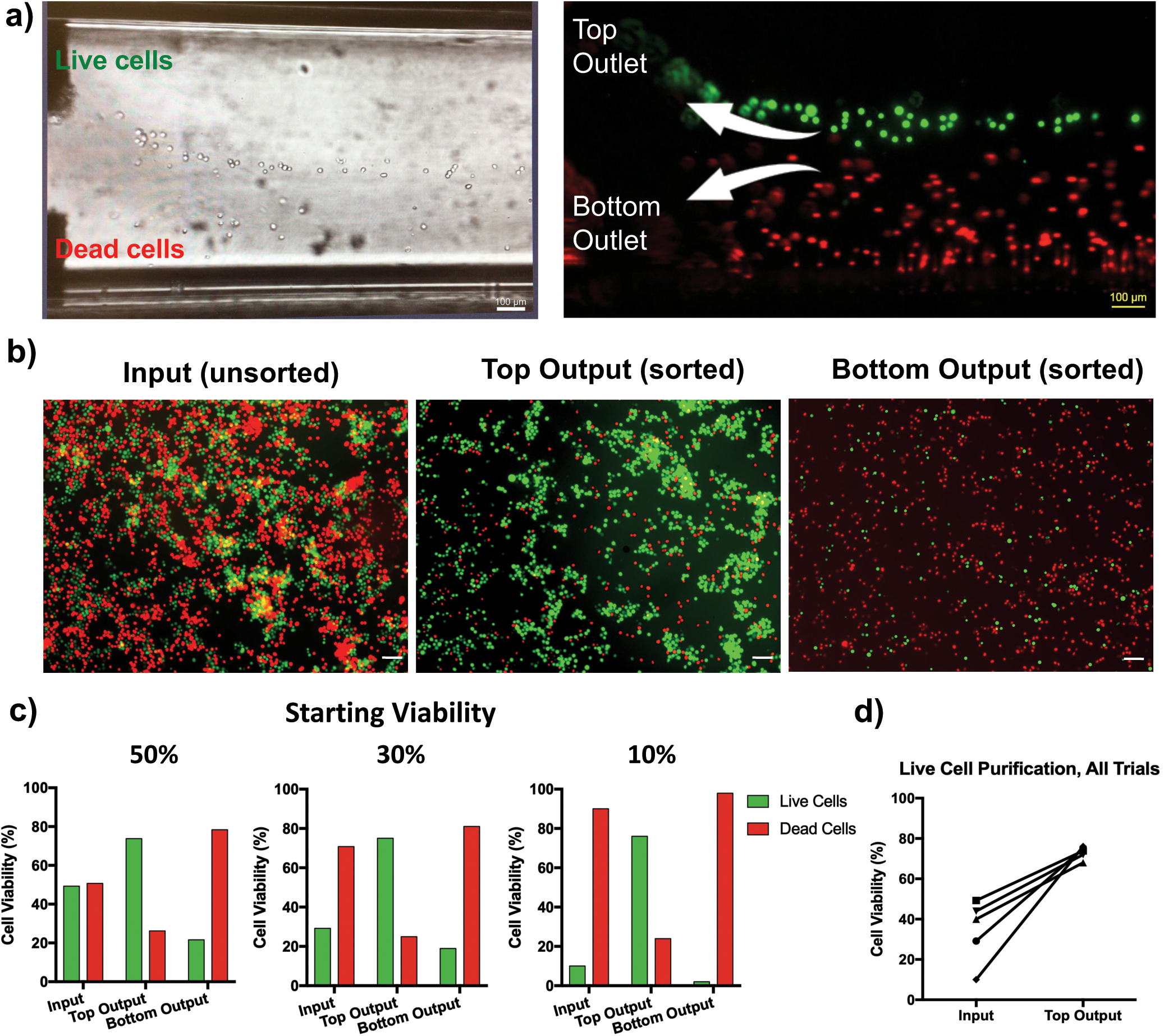
Sorting of live and dead cells via magnetic levitation. **a)** Representative real-time levitation sorting of live and dead cell mixture in 50 mM paramagnetic medium. CelLEVITAS is flexible and versatile. It can sort and enrich live cells from very low input cell concentrations (i.e., 100-500 cells/mL) **(left panel)**, as well as from heterogeneous cell populations from higher input cell concentrations (i.e., >200,000 cells/mL) with higher sample volumes (i.e., 1-10 mL) **(right panel)**. Live and dead cells, despite varied concentrations, separate into distinct collection outlets. **b)** Fluorescent imaging of unsorted cells, purified live cells (green), and purified dead cells (red). The vast majority of cells are sorted into their respective outlet. Imaged cells were counted for calculating separation efficiency. **c, d)** Quantification of purification of live and dead cells. Three trials with different input purities ranging from 10% to 50%, are illustrated in depth; five sorting runs in total were performed. In all the trials, the top outlet achieves high cell viability from as low as 10% input purity, indicating successful sorting and enrichment of live cells, independent of the input purity. Across all trials, magnetic levitational sorting creates a statistically different (p=.0028) population by unequal variance t-test (n=5).

## 3. Discussion

Here, we report the first use of magnetic levitation for the rapid, label-free sorting of viable and non-viable cells within a microfluidic system. The underlying device principles exploit the fact that the cells having undergone cell death are denser than the viable cells, and thus have different levitation characteristics. We utilized these density differences to magnetically levitate, focus, separate, and sort different sub-populations with different cell states. As a model application, we have selectively enriched viable breast cancer cells from heterogeneous mixtures of viable and non-viable sub-populations. Our results show successful sorting from a variety of cell concentrations (100-200,000 cells/mL) and a variety of input purities (10-50%) into consistently high output purities (70-80%). Furthermore, a dramatic removal of dead cells and cell debris was demonstrated. We showed high recovery rates (80-90%) of viable cells with minimal contamination rates of nonviable cells (10-40%). In particular, we demonstrated extremely effective removal of dead cells and debris from samples with extremely low starting cell viability (10%). Thus, CelLEVITAS eliminates the unnecessary centrifugation steps inherent to many current sorting techniques, causing minimal stress on the sorted cells, thereby helping to retain their viability.

This technology has a broad array of uses, spanning a number of diverse applications in biotechnology, regenerative medicine, sequencing, personalized medicine, as well as industrial applications. For instance, the ability to selectively remove inhibitory or toxic dead cells using a simple and rapid procedure results in radically improved cell populations; specifically, regarding viability, antibody/protein yield and consequent functional efficacy improvements. Moreover, given that cell-based therapies are showing promise in treating a variety of diseases, devices capable of gentle, label-free separation of viable cells at high recovery, purity, and throughput could be broadly useful in many areas of biomedicine. This includes the ability to sort cells from limited samples with low viability as demonstrated by our effective enrichment of live cells from a starting sample viability of >10%. Because of its real-time imaging, CelLEVITAS has the capability to sort a wide range of cell concentrations. Here, we demonstrate the sorting from the single cell level up to 200k cells/mL populations of live and dead cell states; however, this microfluidic platform, as previously demonstrated in our point-of-living blood research, can handle more concentrated samples, such as whole red blood sample.^[18]^

In addition to sorting at the extreme ranges of viability and cell count, CelLEVITAS has the advantage of maintaining cell viability post-sorting because of its label-free, density-based approach. Conventional label-based sorting mechanisms, such as fluorescence microscopy, flow cytometry and microplate readers, rely on staining and are prone to high entry costs. Moreover, the label-free nature of CelLEVITAS, as well as the elimination of centrifugation, allows for enhanced sorting of fragile, sensitive cell types as well as clinical samples while maintaining cellular viability. The proposed platform will broadly enable basic research and will overcome the current limitations by improving the viability of the sorted fragile cells as well as increasing the throughput. The following cell types are both large, often fragile and thus can be handled and sorted by the CelLEVITAS technology: macrophages (21 µm),^[19]^ adipocytes (40-300 µm),^[20]^ pancreatic islets (50-500 µm),^[21]^ trophoblast cells (200-300 µm),^[22]^ cardiomyocytes (125×25 µm),^[23]^ model organisms such as *Caenorhabditis elegans* (*C. elegans*) (∼1.15 mm),^[24]^ primary human organoids, tumor spheroids, as well as other large and fragile cells types such as skeletal myocytes, hepatocytes, Schwann cells, and stem cells. In addition, gentle sorting important for sensitive clinical samples with limited sample volume, limited cell concentrations and with low starting viability. This technology saves sorting time by eliminating cell debris and non-target cells. Removing cell debris before sorting increases the efficiency and minimizes the percentage of abort events caused by non-specific influence of the dead cells. Using a clean sample is especially important for the upstream applications of genetic analysis, in particular for sequencing of rare cell populations. Isolating and sequencing only viable cells from the target sample population will dramatically lower the sequencing costs, while increasing the sequencing accuracy and depth.

In future work, CelLEVITAS approach can be explored to profile and predict the drug sensitivity of single cancer cells for clinical applications in precision cancer medicine. Assays that can determine the response of individual tumor cells to cancer therapeutics could greatly aid the selection of drug regimens for personalized treatment of cancer patients. CelLEVITAS can be used to monitor drug responses in real-time, and detect drug-sensitive and drug-resistant sub-populations. Thus, CelLEVITAS is a promising approach for directly assaying single-cell therapeutic responses for precision medicine and personalized medicine.

## 4. Materials and Methods

### CelLEVITAS Levitation Device Design and Flow Setup

The CelLEVITAS levitation device consists of laser-cut pieces of polymethyl methacrylate (PMMA), two permanent neodymium bar magnets (K&J Magnetics) with like poles facing each other and mirrors. For static measurements, a channel (1 mm in height) was placed between the magnets. Angled side mirrors were used for real-time imaging and fine-tuning of levitation profiles of live and dead cell states. Compact device design makes the platform compatible with upright or inverted fluorescence microscopes. For sorting of different cell states, the magnetic levitation-based system was driven by two syringe pumps that withdrew the sample into the sorting system. A heterogeneous sample containing the live and dead/dying cell populations was loaded in a paramagnetic medium and introduced into the inlet. The sample was flowed, levitated and then sorted along the capillary channel, after live and dead/dying cells reached their corresponding equilibrium height within the system. The top outlet was connected to a pump that withdrew the solution containing the live cells (less dense portion), while the bottom outlet was connected to a pump that withdrew the solution containing the dead cells and cell debris (denser portion).

### Cell Culture

MDA-MB-231 breast cancer cells were cultured in Dulbecco’s modified eagle medium (DMEM) (Thermo Fisher) supplemented with 10% FBS and 1% penicillin/streptomycin (Invitrogen Corp.). The cells were grown at 37°C and 5% CO_2_ in a humidified atmosphere.

### System Calibration for Levitation Experiments

Polyethylene beads with different densities (i.e., 1.025 g/mL, 1.030 g/mL, 1.044 g/mL, 1.064 g/mL, 1.089 g/mL) were suspended and levitated in DMEM media with 50 mM paramagnetic medium. Levitation height profiles were imaged and analyzed with in-house developed image analysis software written on MATLAB, as described below.

### Preparation of Live and Dead Cells

MDA-MB-231 cells were centrifuged at 1,200 rpm for 3 minutes at room temperature. Then, cells were re-suspended in DMEM media. Dead cells were prepared by taking a fraction of the total cell suspension (according to the intended final live/dead ratio) and pelleting the cells via centrifugation, followed by re-suspension in PBS and incubation at 60°C for 1 hour. To enable simultaneous observation and confirmation of different cell states under the fluorescent microscope, live cells were stained with calcein (Thermofisher), and dead cells were stained with propidium iodide (PI) (Thermofisher), according to the manufacturer’s instructions.

### General Preparation of Cell Levitation Medium for Experiments

Paramagnetic levitation solutions were prepared with non-cytotoxic, non-ionic, chelated gadolinium ions, in DMEM cell culture media. Gadavist is an FDA-approved, human-injectable, non-toxic magnetic resonance imaging (MRI) contrast reagent.^[25]^ In some cases, calcein and propidium iodide staining was used for biological validation of different cell states.

### Static Levitation of Live and Dead Cells

Live and dead populations of MDA-MB-231 cells were re-suspended in DMEM media with various concentrations of paramagnetic medium (i.e., 30 mM, 50 mM, 100 mM), respectively. For static levitation measurements, 30 µl of this mixture was loaded into the channel and levitated for 30 minutes. Levitation profiles of live and dead/dying cell populations were analyzed and quantified by an in-house developed image analysis software, respectively. In addition, live and dead cell mixture was levitated for 30 minutes and imaged every minute **(Movie S1)**. Cellular positions were monitored to quantify and analyze the levitation trajectories of live and dead cell states.

### Real-Time Levitation Sorting Experiments of Live and Dead Cells in CelLEVITAS Levitation System

Live MDA-MB-231 cells were suspended at 1,200 rpm for 3 minutes. Then, the cell pellet was re-suspended in PBS and cells were stained with calcein for 30 minutes at 37°C. In addition, dead MDA-MB-231 cells were stained with propidium iodide (PI). After staining, dead cells were suspended at 1,200 rpm for 3 minutes and re-suspended in PBS. Then, live and dead cell populations were mixed at various ratios and levitated for 20 minutes at 50 mM paramagnetic medium before sorting. In all sorting experiments, a 2:1 flow ratio was maintained between the top (collecting less dense, live cells) and bottom (collecting denser, dead/dying cells) outlet flow streams **(Movie S2 and S3)**. After sorting, the cells collected from the top and bottom outlets were suspended onto well plates, respectively. Stain-positive cells were detected using the ZenPro2 imaging software (Zeiss). Cells that were calcein-positive (green) were counted as live and cells that were PI-positive (red) were counted as dead/dying. Viable cell recovery was calculated as the number of viable cells in the top collection outlet divided by the sum of viable cells in both the top and bottom collection outlets. Contamination was calculated as the recovery of nonviable cells. Purity, or viability, was calculated as the number of viable cells in the collection outlet divided by the total number of cells eluted via the collection outlet.

### Image Analysis of Levitation Heights of Live and Dead Cells

A MATLAB program for automatic digital image processing was developed. The image datasets were acquired using the ZenPro2 software (Zeiss) and were imported into the code in “.tiff” format. Images were imported as RGB images from the source tiff file and were converted to grayscale. Gaussian smoothing was applied to remove high-frequency noise from the image, followed by a Laplacian operation to filter low-frequency noise. The image was then binarized generating an image that identifies only the cell positive pixels. Pixel values were then summed across rows, and the array was reversed such that 0 corresponds to the bottom of the capillary channel (0 µm) and 1000 (1,000 µm) corresponds to the top of the capillary. These values were summed, with indices counted, to generate a list of heights for all cell-positive pixels. The results were plotted to analyze the distribution of the heights of all cell-containing pixels.

### Statistical analysis

All experiments were repeated three times. Bar graphs are shown as mean ± standard deviation. Unpaired t-test was performed using Excel for statistical analysis. Unpaired t-tests with p-values ≤ 0.05 were considered significant.

## Acknowledgements

N.G.D. acknowledges support from the Career Award at the Scientific Interface (CASI) from the Burroughs Wellcome Foundation (BWF). N.G.D. acknowledges support from the McCormick and Gabilan Faculty Award from Stanford University. Research reported in this publication was also supported in part by the National Cancer Institute (NCI) of the National Institutes of Health (NIH) under award number R25CA217729 (Canary CREST Program). We dedicate this paper to our mentor, Dr. Sam Gambhir.

