## Supplementary File for "CelLEVITAS: Label-free rapid sorting and enrichment of live cells via magnetic levitation"

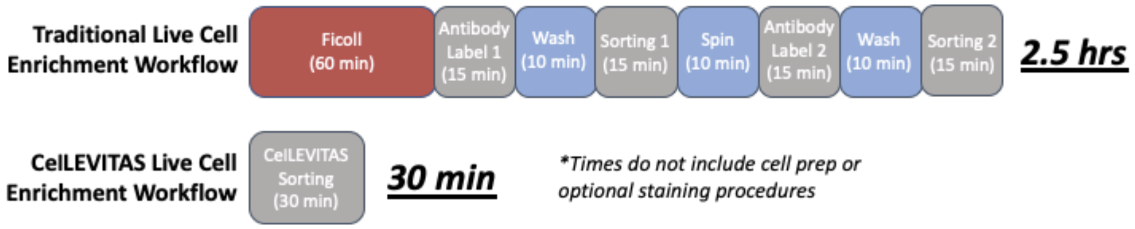


**Supplementary Figure S1: Magnetic Levitation Sorting Accelerates Enrichment of Live Cells Workflow.** The CelLEVITAS procedure reduces the processing time by a factor of five, while maintaining simplicity and avoiding the centrifugation steps. This sorting method is much less time-intensive than traditional methods because it differentiates cells based on inherent density and magnetic properties, rather than surface markers, and thus bypasses the labelling and washing steps.

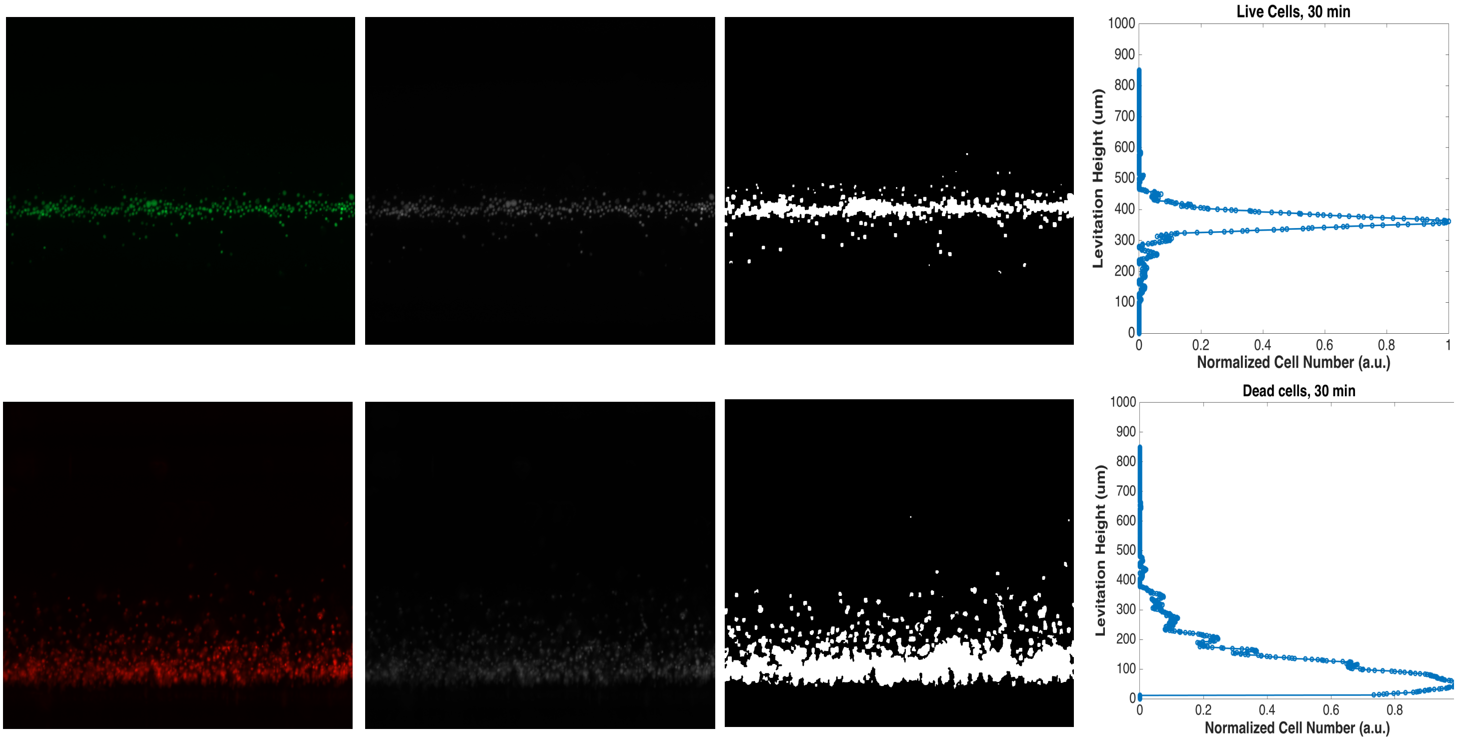


**Supplementary Figure S2. Image analysis of levitation heights of live and dead cells.** A MATLAB program for automatic digital image processing was developed. The image datasets are acquired using the ZenPro2 software (Zeiss) and images with different fluorescent channels (i.e., green representing the live cell levitation band; and red representing the dead/dying cell levitation band) are imported into the code in ".tiff" format. Images were imported as RGB images from the source tiff file and were converted to grayscale. Then, the program uses the magnetic levitation images by examining: (i) if the magnets in the image are tilted, rotates the magnets to make them flat, (ii) between the two magnets by determining the location of the magnets finds the midpoint (coordinate 0), (iii) microparticles determines its location with its radius and (iv) the microparticles found subtracts the distance of the magnet relative to its midpoint. Next, Gaussian smoothing is applied to remove high-frequency noise from the image, followed by a Laplacian operation to filter low-frequency noise. The image is then binarized generating an image that identifies only the cell positive pixels. Pixel values are then summed across rows, and the array is reversed such that 0 corresponds to the bottom of the capillary channel (0 μm) and 1000 (1,000 μm) corresponds to the top of the capillary. These values are summed, with indices counted, to generate a list of heights for all cell-positive pixels. The results are plotted to analyze the distribution of the heights (y-axis, levitation height) of all cell-containing pixels (x-axis, normalized cell number (a.u.)).

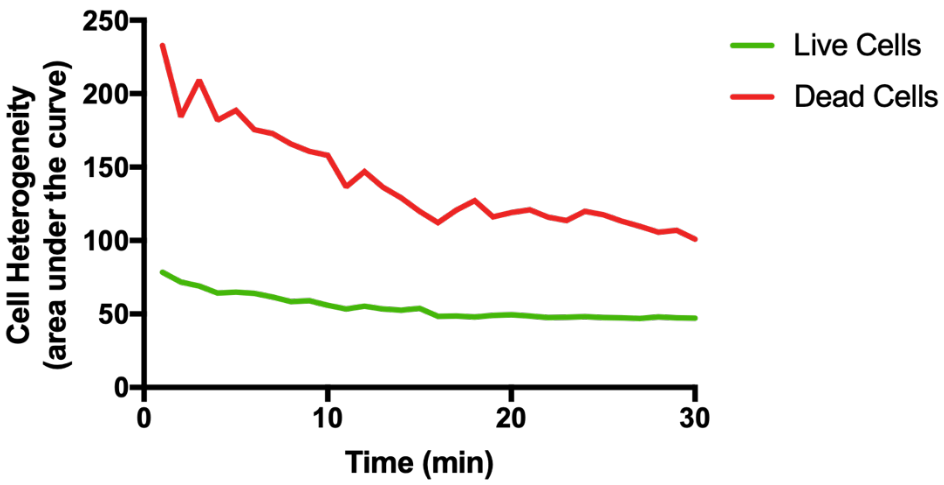


**Supplementary Figure S3. Heterogeneity of live and dead cell populations over time.** Population heterogeneity was calculated as the integral of the normalized cell number of a given cell population. This is equal to the area under the curves graphed in Figure 3c. As the cell number is normalized, this analysis on the statistical distribution of cell heterogeneity considers only how spread a cell population is across many levitation planes. Larger area under the curve indicates less clustering of cells around a mean levitation height. Declining live and dead cell heterogeneities guides the sorting parameters and justify that the optimal static levitation period before sorting.

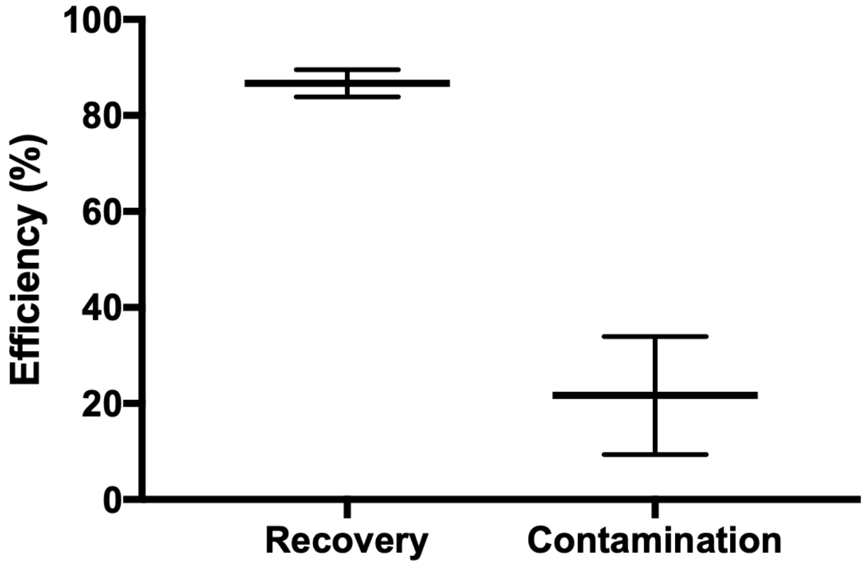


**Supplementary Figure S4. Viable cell recovery and nonviable cell contamination when sorting.** The top outlet consistently recovered 80-90% of all viable cells sorted, while only recovering under 30% of all nonviable cells. Thus, CelLEVITAS achieves high purity in the top outlet without sacrificing recovery rate. The contamination rate has greater variability, likely due to the heterogeneity in levitation height of dead cells. Recovery and contamination data are graphed only for the optimal flow rate of 1.2 mL/hr with a 2:1 top: bottom ratio.

**Supplementary Table S1. Comparison of measured densities of live and dead cell states with the literature.**

| | Density (g/mL) | |
|---|---|---|
| | **Measured (mean ± s.d.)** | **Literature (range)** |
| Viable MDA-MB-231 cells | 1.043 ± 0.002 | 1.044 ± 0.018 (Ref) |
| Non-viable MDA-MB-231 cells | 1.075 ± 0.004 | - |

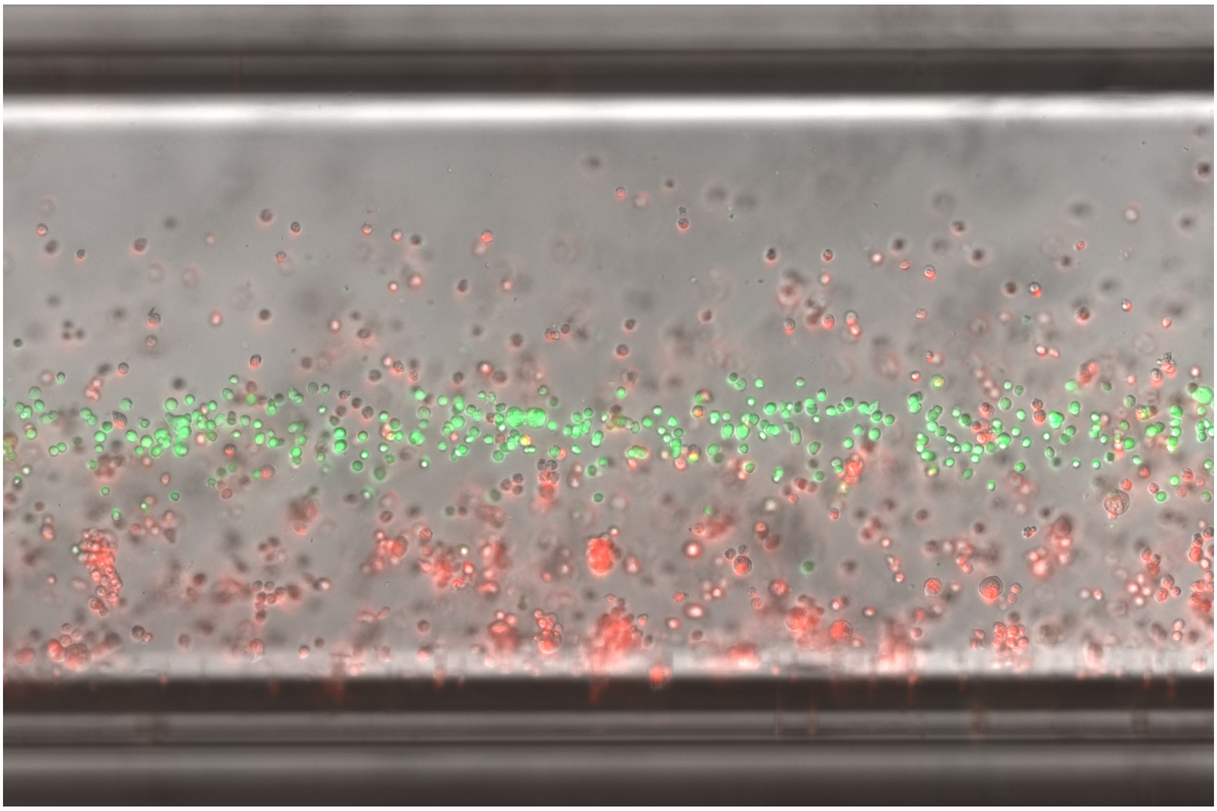

**Supplementary Movie S1.** Static levitation and magnetic focusing of heterogonous mixture of live and dead cells at 50 mM paramagnetic medium. Live and dead cell mixture was levitated for 30 minutes and imaged every minute. Cellular positions were monitored and the levitation trajectories of live and dead cell states were analyzed.

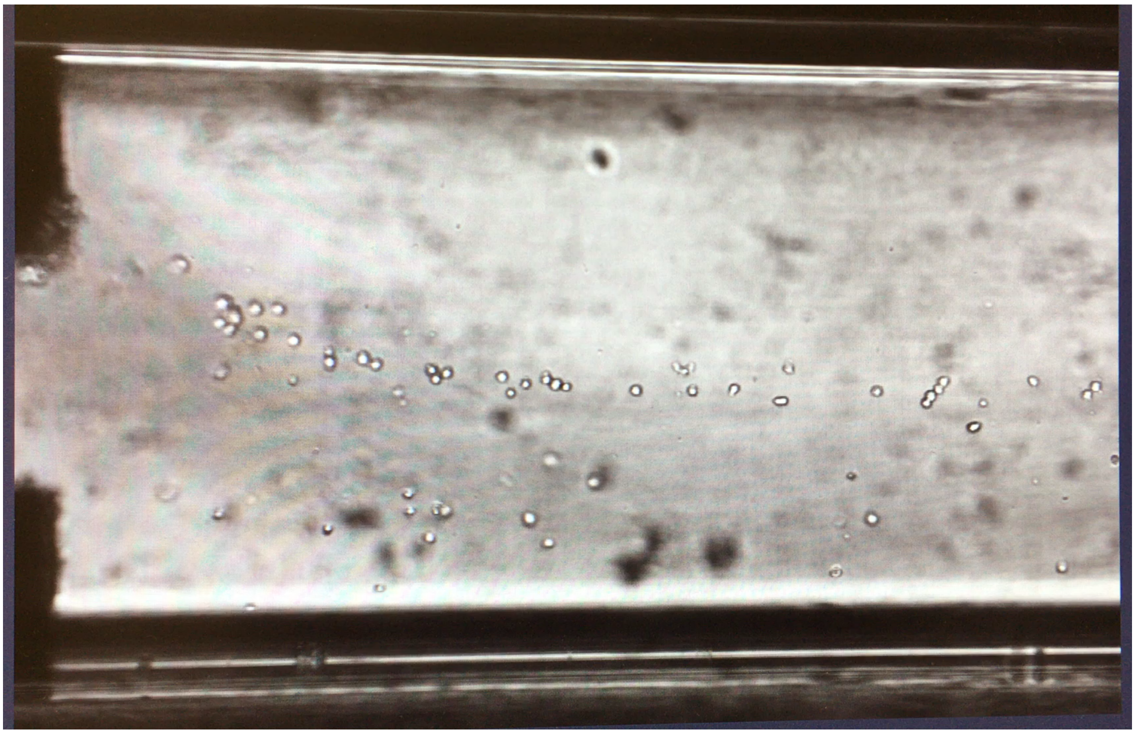

**Supplementary Movie S2.** Rapid magnetic sorting and enrichment of live cells from very low number or input cell concentrations (i.e., 100-500 cells).

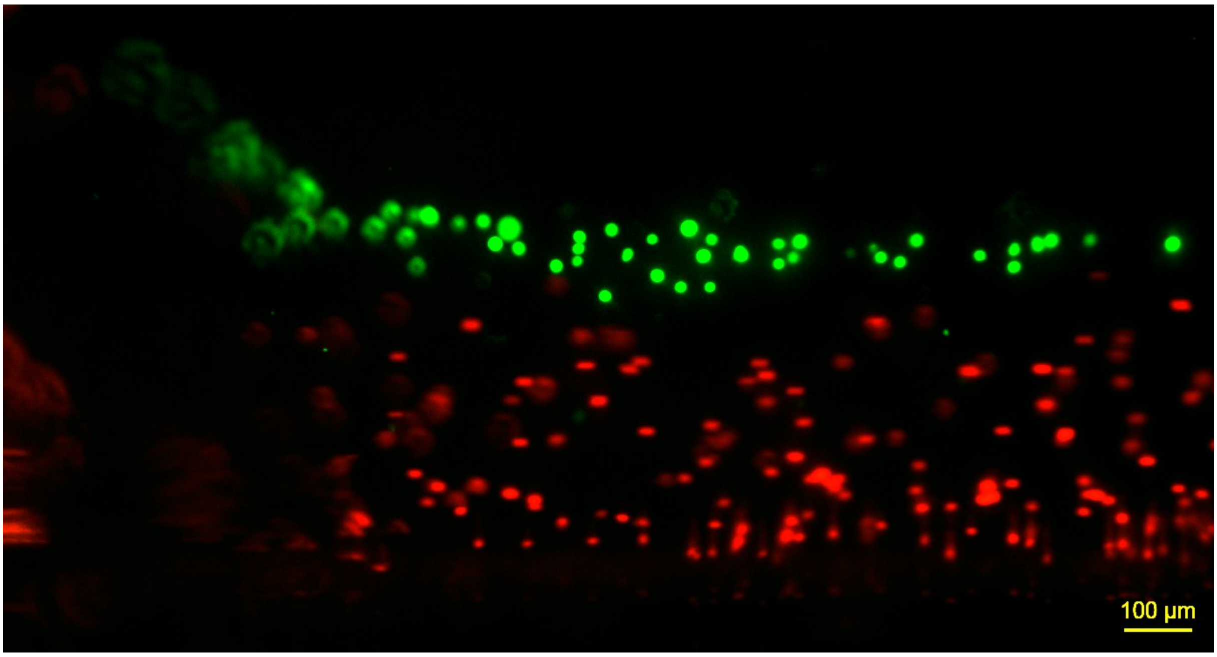


**Supplementary Movie S3.** Rapid magnetic sorting and enrichment of live cells from heterogeneous cell populations from higher input cell concentrations (i.e., >200,000 cells/mL).
